# Natural reversal of cavefish heart asymmetry is controlled by Sonic Hedgehog effects on the left-right organizer

**DOI:** 10.1101/2023.12.13.571559

**Authors:** Mandy Ng, Li Ma, Janet Shi, William R. Jeffery

## Abstract

The direction of left-right (L-R) visceral asymmetry is conserved in vertebrates. Deviations of the asymmetric pattern are rare, and the underlying mechanisms are not understood. Here we use the teleost *Astyanax mexicanus*, a single species consisting of surface fish with normal left-oriented heart asymmetry and cavefish with high levels of reversed right-oriented heart asymmetry, to explore natural changes in L-R determination. We show that Sonic Hedgehog (Shh) signaling is increased at the posterior midline, Kupffers Vesicle (KV), the teleost L-R Organizer (LRO), is enlarged and contains longer cilia, and the number of dorsal forerunner cells, the KV precursors, is increased in cavefish. Furthermore, Shh overexpression in surface fish induces KV modifications and changes in heart asymmetry resembling the cavefish phenotype. Asymmetric expression of the Nodal antagonist *dand5* is equalized or reversed in the cavefish KV, and Shh overexpression in surface fish mimics changes in cavefish L-R *dand5* asymmetry and shifts Nodal laterality in the lateral plate mesoderm. Our results show that naturally occurring modifications in cavefish heart asymmetry are controlled by the effects of enhanced Shh signaling on LRO function.

**Summary:** Natural reversal of cavefish heart asymmetry is controlled by enhanced Sonic Hedgehog signaling at the posterior midline, which modifies left-right organizer function and subsequent Nodal laterality.

## Introduction

Bilaterally symmetric animals typically show mirror image symmetry of external organs, such as the eyes and limbs, but also exhibit remarkable asymmetries of internal organs (Blum and Ott, 2018; Blum et al., 2014). In vertebrates, the position of the heart is biased toward the left side of the plane of bilateral symmetry (midline), whereas the liver is mostly located on the right side of the body cavity (Grimes and Burdine, 2017). This pattern of left-right (L-R) visceral asymmetry is conserved among diverse vertebrate species, although low levels of partial or complete inversions have been detected in some natural populations (Malicki et al., 2011; Wang et al., 2011; Noël et al., 2013) or can be induced by mutations in genes governing L-R determination (Chen et al., 2001; Tan et al., 2007).

L-R visceral asymmetry is established during vertebrate embryogenesis through the activity of the L-R Organizer (LRO). In amphibians, fish, and some mammals, the LRO is ciliated, and leftward ciliary beat and fluid flow determine the pattern of visceral organ asymmetry (Blum et al., 2014; Hamada, 2020). The teleost LRO, known as Kupffer’s vesicle (KV), is a spherical structure lined by motile cilia (Essner, et al, 2005; Kramer-Zucker et al., 2005; Bajoghli et al., 2007; Sampaio et al., 2014; Gokey et al., 2016). The KV develops from the dorsal forerunner cells (DFC), derivatives of the Wilson cells (Warga and Kane, 2018), which converge toward the midline at the leading edge of the blastoderm during embryonic shield formation (Cooper and D’Amico, 1996; Oteiza et al., 2008). Leftward fluid flow reduces expression of the Cerberus/Dan family gene *dand5*, encoding a Nodal antagonist, on the left side but not the right side of the KV (Hashimoto et al., 2004; Smith and Uribe, 2021), which in turn promotes expression of the Nodal family gene *spaw* in the left lateral plate mesoderm (LPM) (Long et al., 2003). Spaw subsequently activates the downstream homeobox gene *pitx2*, and the Spaw-Pitx2 signal is propagated from posterior to anterior in LPM on the left side of the embryo, which determines the L-R pattern of visceral organ asymmetry (Grimes and Burdine, 2017). Spaw is prevented from spreading across the midline by a barrier consisting of BMP4, the Nodal antagonist Lefty1, which is expressed in the notochord, and the Nodal antagonist Lefty2, which is expressed in the left LPM (Bisgrove et al, 1999; Chocron et al., 2007; Lenhardt et al., 2011; Smith et al. 2011). A similar chain of events is initiated in mouse embryos by leftward flow fluid from the LRO resulting in Nodal-Pitx2 activation exclusively in the left LPM, suggesting a conserved sequence of L-R asymmetry determining events in vertebrates with ciliated LROs (Montague et al., 2018; Hamada, 2020; Skenker-Ravi et al., 2022).

Although the mechanisms responsible for establishing L-R visceral asymmetry in vertebrates are fairly well known (Smith and Uribe, 2021), those controlling natural changes in asymmetric patterning are not understood. One of the reasons for this gap in knowledge is that abnormal asymmetry is rare and difficult to study in most vertebrates. For example, heart asymmetry is inverted in only about 1 of 5,000 to 7,000 human births (Shiraishi and Ichikawa, 2012). Here we capitalize on the teleost *Astyanax mexicanus* (Jeffery, 2001; 2020), which provides a unique system for investigating the molecular and evolutionary mechanisms underlying modifications in L-R determination (Ma et al., 2021a).

*Astyanax mexicanus* is a single species with a surface-dwelling form (surface fish) and a cave-dwelling form (cavefish) (Jeffery, 2001; 2020). Cavefish have evolved regressive and constructive traits that differ from surface fish, including eye degeneration, loss of pigmentation, increased olfactory and taste sensitivity, larger mouthparts and fat deposits, and changes in feeding and social behaviors (Jeffery, 2001; 2020; Xiong et al., 2018; Kowalko, 2020). Many of these traits are accompanied by enhanced embryonic midline structures, namely the pre-chordal plate, notochord, and floor plate of the neural tube, highlighted by overexpression of the Sonic Hedgehog (Shh) signaling system (Yamamoto et al., 2004; 2009; Menuet, et al. 2007; Hinaux et al., 2016; Ren et al., 2018; Sifuentes-Romero et al., 2020). Modifications in L-R asymmetry have also evolved in cavefish: whereas surface fish, like zebrafish and other vertebrates, show heart asymmetry biased toward the left side of the midline, some cavefish populations exhibit up to 20-30% right-oriented heart asymmetry (Ma et al., 2021a). Reciprocal hybridization between surface fish males with normal heart asymmetry and cavefish females with reversed heart asymmetry has shown that a significant number of the F1 hybrid progeny exhibit the inverted heart asymmetry phenotype of their mothers, indicating that maternal genetic effects are involved in cavefish L-R asymmetry (Ma et al., 2021a). The laterality of Spaw-Pitx2 signaling in the LPM is also modified in cavefish (Ma et al., 2021a), suggesting that maternal factors may control L-R asymmetry through unknown zygotic events acting upstream of Nodal laterality. The purpose of the present investigation was to identify these zygotic factors and understand how they influence L-R asymmetry in cavefish.

We show that changes in cavefish heart asymmetry are controlled by upregulation of the Shh pathway at the posterior midline, in a region we term the post-chordal plate. Shh overexpression in the post-chordal plate affects DFC number, KV size and ciliation, L-R *dand5* asymmetry, Spaw-Pitx2 laterality in the LPM, and reverses the pattern of L-R heart asymmetry. The results further suggest that the evolution of reversed L-R visceral asymmetry in cavefish is linked to the development of constructive and regressive traits controlled by the Shh midline signaling system.

## Results

### The left-right organizer is modified in cavefish embryos

To investigate the possibility of differences between the surface fish and cavefish LROs, we compared KV sizes and ciliation between surface fish families with >95% D-looping cardiac tubes (SF-D), cavefish families with 90-95% D-looping cardiac tubes (CF-D), and cavefish families with 20-30% L-looping cardiac tubes (CF-L/D) (Figs. 1 and 2). KVs were measured at four consecutive stages during embryonic segmentation: the 10-12 somite stage, the 13-15 somite stage, the 16-18 somite stage, and the 19-21 somite stage (Fig. 1A-I, N). The mean KV area of CF-L/D embryos was significantly larger than SF-D and CF-D embryos between the 10 to 15 somite stages, but KV sizes were similar in surface fish and both cavefish families from the 16 to the 21 somite stages (Fig. 1A-I, N). Thus, as previously reported in zebrafish (Gokey et al., 2016), *Astyanax* KVs initially inflate and then gradually deflate during somitogenesis. However, most CF-L/D KVs inflated more rapidly and to a larger size than SF-D or CF-D KVs. Surprisingly, a small subset of CF-D and CF-L/D embryos exhibited multiple KVs: two or three KVs of similar size were arranged in a row at the ends of a single, posteriorly split notochord (Fig. 1J, M; also see Fig. 4M), which was not seen in SF-D (Fig. 1M) or to our knowledge in the LROs of any other vertebrate species (Blum et al., 2009). The multiple cavefish KVs were confirmed in CF-L/D embryos by in situ hybridization with the KV marker gene *c1orf127* (Fig. 1K, L). The results show that LRO size and number are modified between surface fish and cavefish, and that these differences are especially pronounced in CF-L/D embryos.

**Figure 1.**
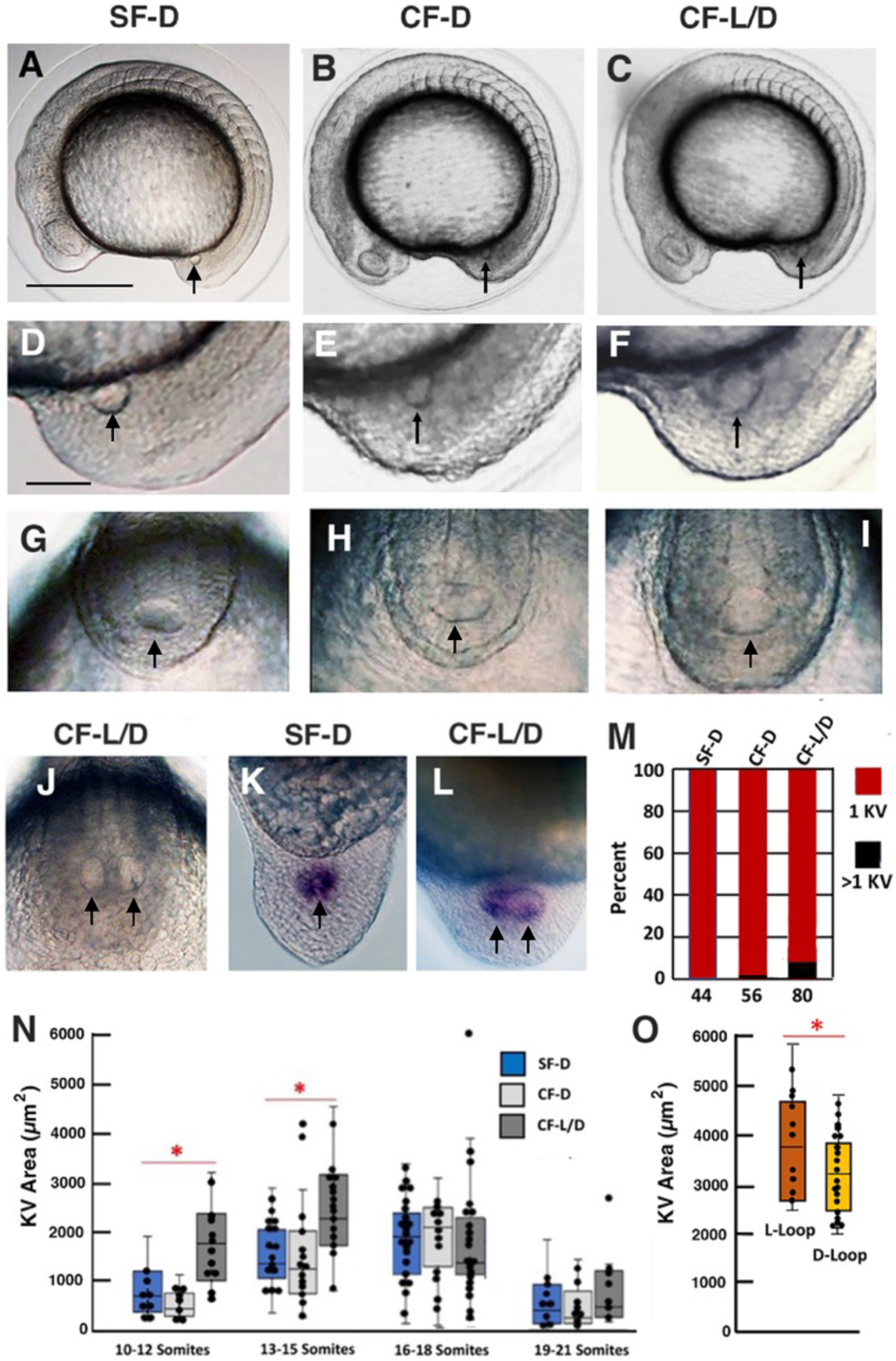
Left-Right Organizer modifications and relationship to heart asymmetry in cavefish. A-C. SF-D (A), CF-D (B), and CF-L/D (C) embryos at the 13-15 somite stage. Scale bar in A: 100 µm; magnification is the same in A-C. D-I: Tailbud regions of 15 somite SF-D (D, G), CF-D (E, H), and CF-L/D (F, I) embryos viewed from the lateral (D-F) and dorsal (G-I) sides. J. A CF-L/D embryo showing two KVs and a posteriorly split notochord. K, L. SF-D (K) and CF-L/D (L) 15 somite embryos stained with the KV marker gene *c1orf127*. L. Scale bar in D: 30 µm; magnification is the same in D-L. Arrows: KVs. M. Bar graph showing the percentage of 15-somite SF-D, CF-D, and CF-L/D embryos with multiple KVs. The number of embryos is shown at the base of each bar. N. KV size differences in SF-D, CF-D, and CF-L/D during the 10-12, 13-15, 16-18, and 19-21 somite stages. Asterisks: Significance at p < .00001 and p = .00141 from left to right. O. The relationship between KV size and the direction of cardiac tube looping heart in CF-L/D. Asterisk: Significance at p = .016018. Statistical analysis by one-way ANOVA with Tukey HSD.

**Figure 2.**
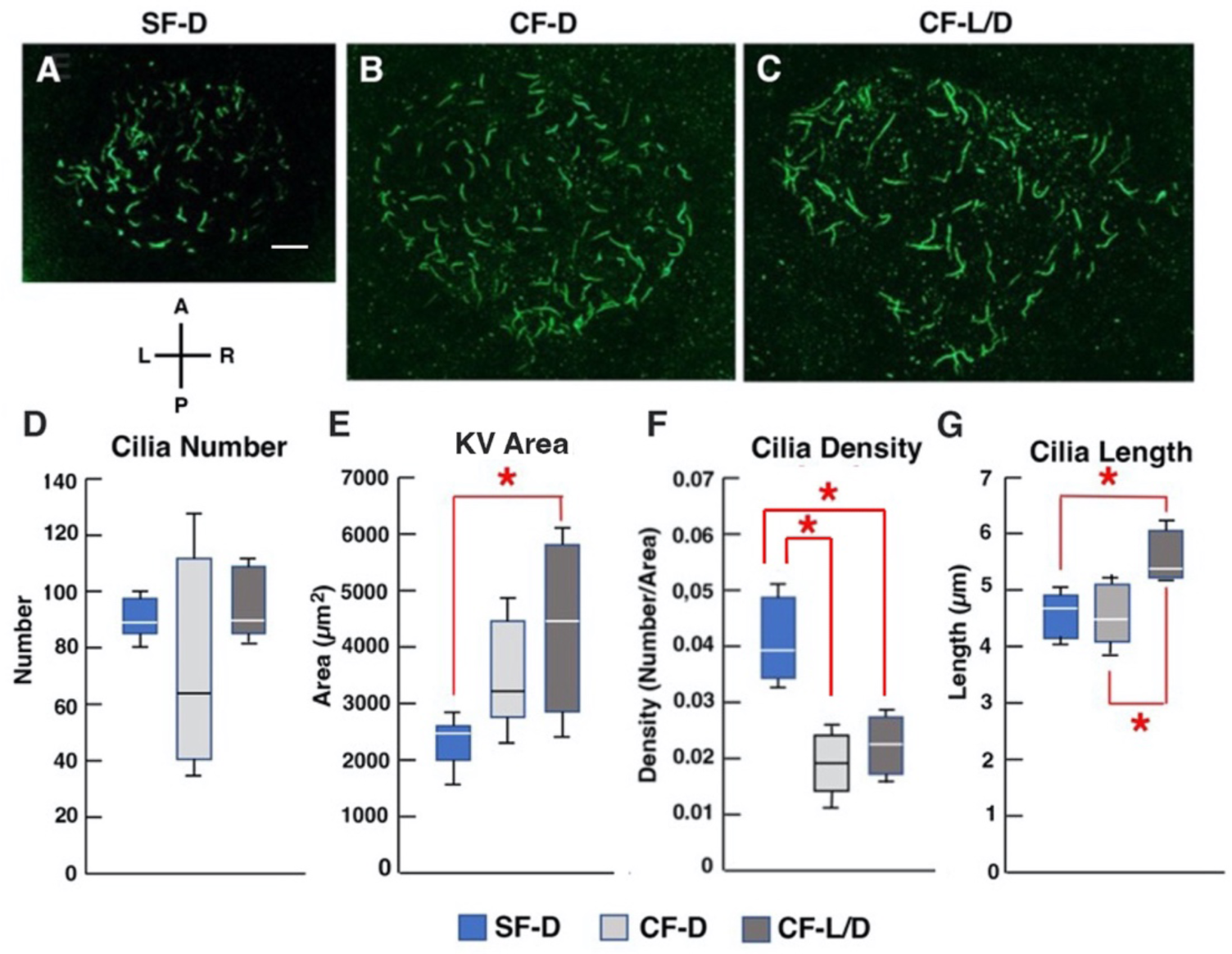
KV ciliation in surface fish and cavefish embryos. A-C. SF-D (A), CF-D (B), and CF-L/D (C) KVs stained with alpha-acetylated tubulin antibody at the 15 somite stage. Scale bar in A: 10 µm: magnification is the same in A-C. D-F. Box and whisker plots showing the quantification of KV cilia number (D), area (E), density (F), and length (G) in 15 somite stage SF-D, CF-D, and CF-L/D embryos. Horizontal lines in boxes: Medians. Vertical lines: whiskers representing the 25^th^ percentile. N = 9 (SF-D), 5 (CF-D), and 5 (CF L-D). Asterisks: significances in (E) p = .00667, (F), p = .0075 (upper) and p = .00254 (lower), and (G) p = .00262 (upper) and p = .00303 (lower). Statistics by One way ANOVA with Tukey HSD.

To determine whether surface fish and cavefish KVs differ in ciliation, SF-D, CF-D, and CF-L/D embryos were stained with alpha-acetylated tubulin antibody at the 13-15 somite stage, and cilia number, density, and length were quantified (Fig. 2A-C). A significant increase the length of cilia was found in CF-L/D KVs relative to CF-D and SF-D KVs (Fig. 2F). No significant differences were found in the number of KV cilia between SF-D, CF-D, and CF-L/D (Fig. 2D). However, due to larger KV areas, the density of cilia was significantly reduced in the KVs of CF-D and CF-L/D embryos compared to SF-D embryos (Fig. 2E, F). Taken together, the results indicate that the KV is modified in cavefish relative to surface fish, and that KV modifications are more extreme in CF-L/D embryos.

The relationship between KV modifications and the direction of heart asymmetry was determined by measuring the KV sizes of individual 13-15 somite CF-L/D embryos, then raising each embryo separately and assaying for cardiac tube looping at about 2.5 days post-fertilization (dpf). The results showed that CF-L/D embryos with L-looping heart tubes originated mostly from embryos with significantly larger KVs than CF-L/D embryos with D-looping cardiac tubes (Fig. 1O).

### Dorsal forerunner cells and *foxj1a* expression are increased in cavefish embryos

We next sought to understand the causes of differences between the surface fish and cavefish KVs. The KV is formed by coalescence of DFC along the dorsal midline at the shield stage (Cooper and D’Amico, 1996; Oteíza et al., 2008). The DFC of SF-D, CF-D, and CF L/D embryos were stained with Syto 11 and quantified at 70%-80% epiboly (Fig. 3A-I). As in zebrafish (Moreno-Ayala et al., 2021), DFC numbers varied greatly within *Astyanax* SF-D, CF-D, and CF-L/D embryos (Fig 3I). However, the mean number of DFC was significantly larger in CF-L/D embryos relative to CF-D and SF-D embryos (Fig. 3I). The DFC of SF-D (Fig. 3A, B), CF-D (Fig. 3C, D), and most CF-L/D embryos (Fig. 3 E, G) coalesced into a single focal point during gastrulation, whereas a subset of CF-L/D embryos DFC formed several focal points (Fig. 3F, H), a likely explanation for the development of multiple KVs (Fig. 1K, L). The results suggest that the larger and multiple KVs of CF-L/D embryos can be explained by the development of more DFC.

**Figure 3.**
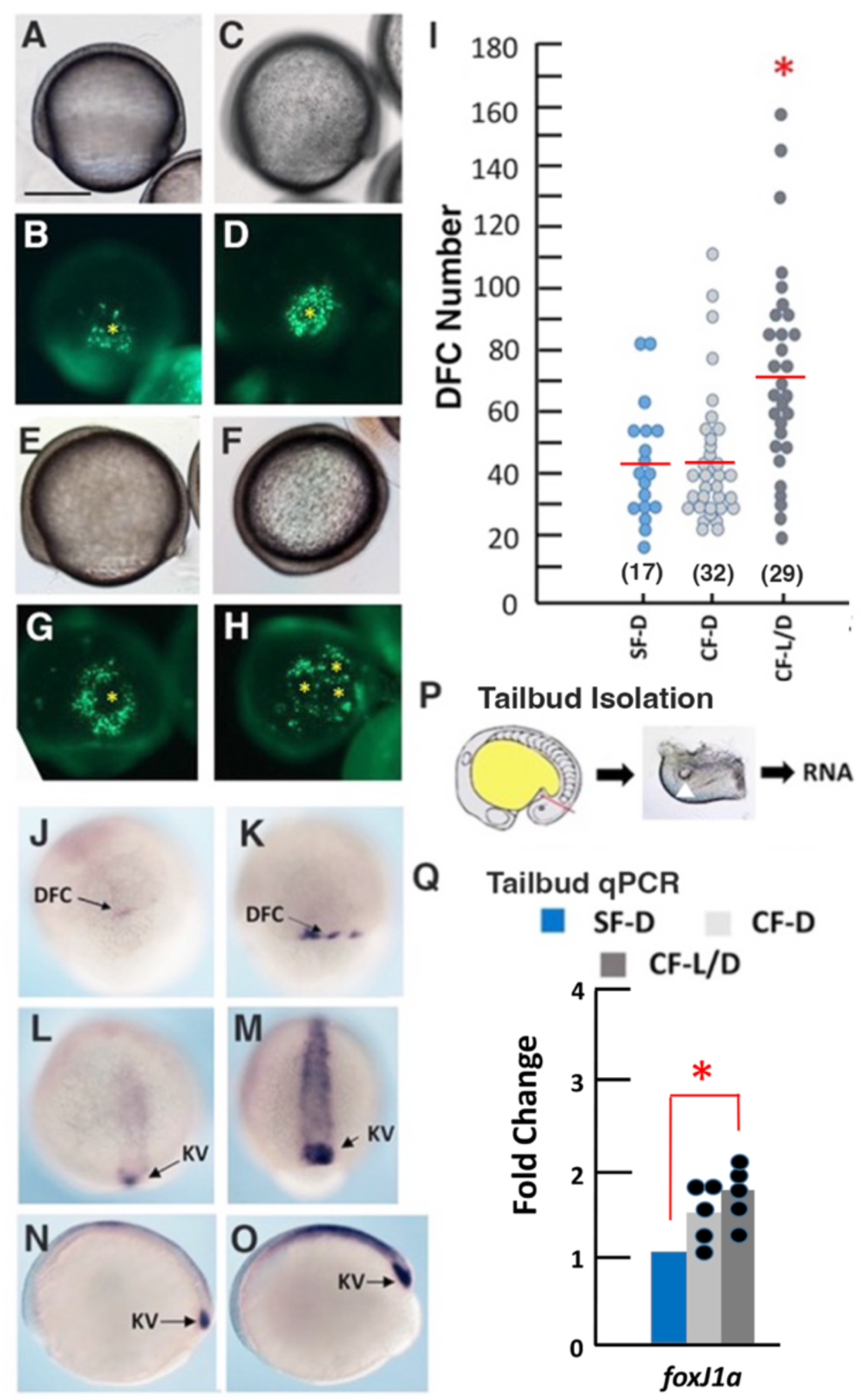
Increase in dorsal forerunner cells and *foxj1a* expression in cavefish embryos. A-H. Dorsal forerunner cells (DFC) at 70-80% epiboly in SF-D (A, B), CF-D (C, D), and CF-L/D (E-H). A, C, E, F: bright field images. B, D, G, H: Syto 11 stained fluorescence images. Asterisks indicate the foci of DFC coalescence. I. DFC quantification in SF-D, CF-D, and CF-L/D embryos at 70-80% epiboly. Red horizontal lines: Means. Number of embryos quantified is shown in parentheses. Asterisk: significance at p = .000102. Statistics by one-way ANOVA with Tukey HSD. J-O. In situ hybridization showing *foxj1a* expression in SF-D (J, L, N) and CF-L/D (K, M, O) embryos at the 50% epiboly (J, K), 70% epiboly (L, M) and early tailbud (N, O) stages. Scale bar in A: 100 µm: magnification is the same in A-H and J-O. P. Diagram depicting the isolation of surface fish and cavefish tailbuds for RNA extraction in tailbud qRT-PCR quantifications. Diagonal red line: position of tailbud amputation. Center: Image of isolated cavefish tailbud with KV (arrowhead). Q. Tailbud qRT-PCR quantification of *foxj1a* expression in SF-D, CF-D, and CF-L/D embryos at the 13-15 somite stage. Asterisk: significance at p = .006802. Statistics by one-way ANOVA with Tukey HSD.

The DFC, and subsequently the KV, express *foxj1a,* a master regulator of motile cilia development (Yu et al., 2008; Tavares et al., 2017), along the midline during gastrulation and axis development. Therefore, we conducted in situ hybridization and quantitative reverse transcriptase polymerase chain reaction (qRT-PCR) to determine whether *foxj1a* expression is changed in cavefish embryos (Fig. 3J-O). In situ hybridization showed that *foxj1a* staining is stronger in the DFC of CF-L/D than SF-D embryos during gastrulation (Fig. 3J, K) and along the midline and in the KV during the 13-15 somite stage (Fig. 3L-O). To quantify *foxj1a* expression specifically in the KV region, we developed an approach in which qRT-PCR was performed with RNA extracted from tailbuds surgically removed from 13-15 somite stage surface fish and cavefish embryos (Fig. 3P), and this tailbud qRT-PCR approach was also used in subsequent parts of this study (see below). Tailbud qRT-PCR quantification showed significant increases in *foxj1a* mRNA levels in CF-D and CF-L/D embryos compared to SF-D embryos (Fig. 3Q). The results suggest that increases in *foxj1a* expression and the number of DFC may account for structural modifications in the cavefish LRO.

### Shh and BMP signaling is increased at the posterior midline of cavefish embryos

The *foxj1a* gene is a direct target of Shh signaling (Cruz et al., 2010). Furthermore, previous studies showed that Shh signaling is expanded in the pre-chordal plate at the anterior midline of cavefish compared to surface fish embryos and that this increase subsequently contributes to lens apoptosis, eye degeneration, and other phenotypic changes in cavefish (Yamamoto et al., 2004, 2009; Menuet et al., 2007). Accordingly, we hypothesized that increased Shh signaling could also be responsible for excessive KV development at the posterior midline of cavefish embryos. To investigate this possibility, the expression of *shh* and its autoregulated receptor *ptch2* were compared in the KV region of 13-15 somite surface fish and cavefish embryos by in situ hybridization and qRT-PCR (Fig. 4). In addition to overexpression at the anterior midline, we found that *shh* and *ptch2* expression are expanded laterally along the posterior midline of cavefish embryos (Fig. 4A-L, Q), suggesting increased Shh signaling in the latter region. Furthermore, when two KVs were present in CF-L/D embryos, *shh* expression was associated with a spur of the notochord extending to each of them (Fig. 4M), and expanded *ptch2* expression surrounded both KVs (Fig. 4N) . Tailbud qPCR (Fig. 3P) confirmed the increase in *shh* expression at the posterior midline in CF-L/D relative to SF-D embryos (Fig 4R).

**Figure 4.**
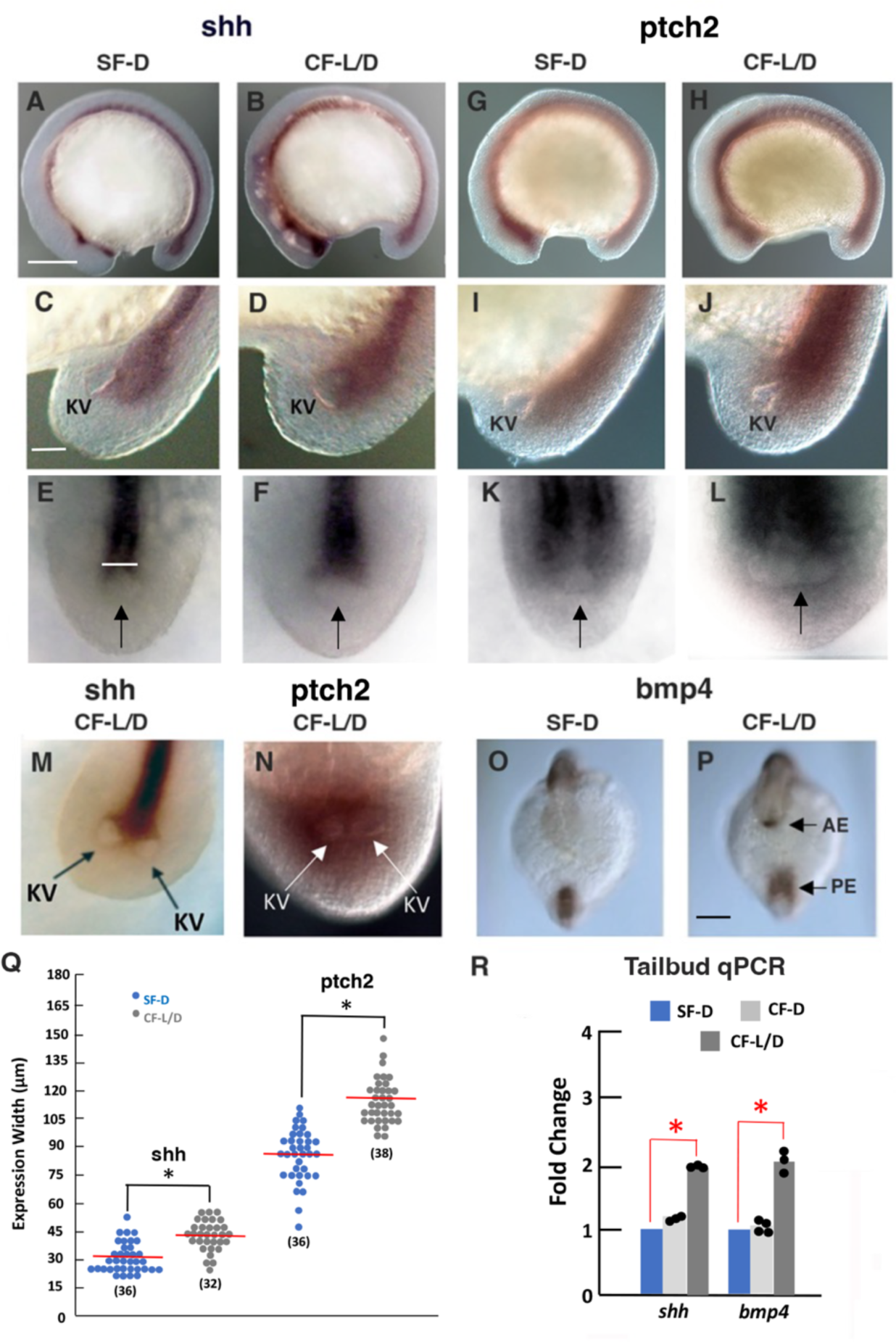
Shh and BMP signaling is increased along the posterior midline in cavefish embryos. A-L. In situ hybridization showing *shh* (A-F) and *ptch2* (G-L) expression in 13-15 somite SF-D (A, C, E, G, I, K) and CF-L/D (B, D, F, H, J, L) embryos. Lateral views of embryos (A, B, C, D) and tailbuds (C, D, I, J) with anterior poles on the left. E, F, K, L. Dorsal views of tailbuds with anterior poles at the top. Vertical line in E: width of expression measurement area (see Q). Arrows in E, F, K, L. Position of KV (out of focus). M, N. In situ hybridization showing *shh* (M) and *ptch2* (N) expression in CF-L/D embryos with two KVs. Scale bar in A is 50 µm; same magnification in A, B, G, H. Scale bar in C is 30 µM: same magnification in C-N. O, P. In situ hybridization showing *bmp4* expression in 13-15 somite SF-D and CF-L/D embryos. Scale bar in P is 50 µm; same magnification in O, P. AE: anterior expression domain. PE: posterior expression domain. Q. Widths of *shh* and *ptch2* expression domains in the KV region of SF/D and CF-L/D embryos. Red horizontal lines: means. Number of embryos quantified is shown parentheses. Asterisks: significance at p < .00001. Statistics by one-way ANOVA with Tukey HSD. R. Tailbud qRT-PCR quantification (see Fig. 3P, top) of *shh* and *bmp4* mRNA in 13-15 somite SF-D, CF-D, and CF-L/D embryos. Asterisks: significance at p < .00001 and p = .000533 from left to right. Statistics by one-way ANOVA with Tukey HSP.

Along with Shh signaling, BMP4 signaling is also expanded in the pre-chordal plate of cavefish embryos (Pottin et al., 2011; Hinaux et al., 2016). Therefore, *bmp4* expression was compared in 13-15 somite surface fish and cavefish embryos. In situ hybridization showed that *bmp4* expression appears in the pre-chordal plate, as described previously, and is also expanded along the posterior midline surrounding the KV in CF-L/D embryos compared to SF-D embryos (Fig. 4O, P), and an increase in *bmp4* expression was confirmed by tailbud qRT-PCR in CF-L/D relative to SF-D embryos (Fig. 4R). It is concluded that Shh and BMP4 signaling are enhanced in a posterior signaling center surrounding the cavefish KV.

### Shh overexpression affects the left-right organizer, eye development, and heart asymmetry

To determine the effects of increased Shh signaling on KV development and heart asymmetry, we treated SF-D embryos with the Smoothened agonist SAG, which has been effective in specifically increasing Shh signaling in different vertebrate systems (Chen et al., 2002; Lewis and Krieg, 2014; Negretti et al., 2022). In these experiments, SAG or DMSO (control) treatment began at about 50% epiboly and lasted until the 13-15 somite stage (8 hours), when aliquots of control and SAG-treated embryos were removed for KV measurements, KV cilia staining, and mRNA quantification (Fig. 5A). The remaining embryos were then raised to later stages to confirm the effects of Shh overexpression on cavefish development (Yamamoto et al., 2004; 2009): aliquots were removed at 2.5 dpf for investigating lens apoptosis and eye primordium size and at 3 months post-fertilization to determine the long-term effects on eye development (Fig. 5A). qRT-PCR showed that SAG treatment significantly increased *ptch1*, *gli1*, and *nkx2.1* mRNA levels relative to DMSO controls (Fig. 5B), indicative of enhanced Shh signaling. Furthermore, the lens showed apoptosis (Fig. 5C, D), the ventral side of optic primordia was reduced (Fig. 5E-G), and fry developed extremely regressed eyes (Fig. 5H, I) in SAG-treated but not control SF-D embryos. Together, these results suggest that SAG is effective in increasing Shh signaling and producing eye degenerative phenotypes resembling cavefish in surface fish.

**Figure 5.**
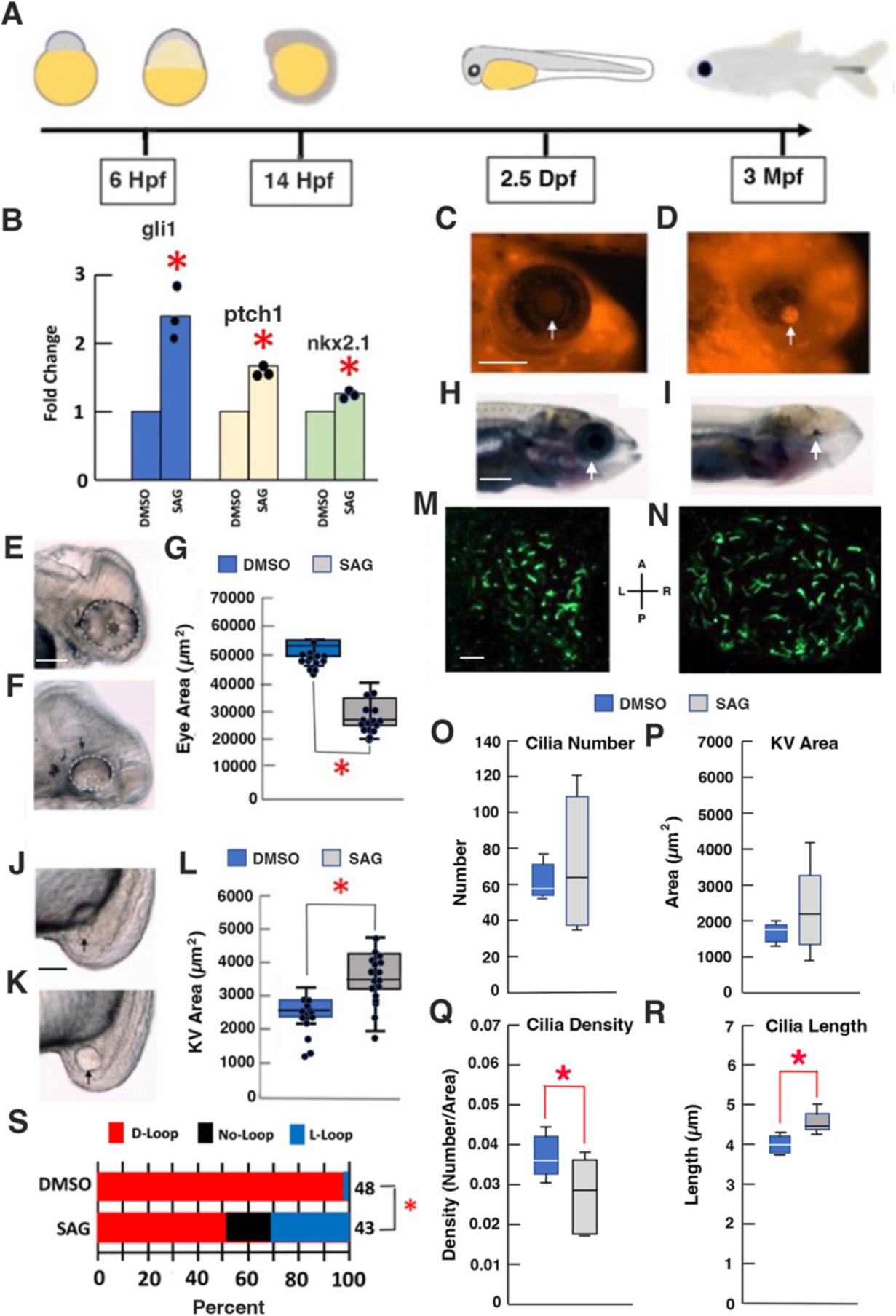
Effects of Sonic Hedgehog overexpression on KV size and ciliation, eye development, and cardiac looping asymmetry in surface fish. A. The timeline of Shh overexpression experiments. SAG or DMSO (control) were added to SF-D embryos at 6 hrs post-fertilization (50% epiboly). After washing into water without SAG or DMSO at 14 hrs post-fertilization (13-15 somite stage, aliquots were removed for KV size determination, RNA was isolated for qPCR, aliquots were removed at 2.5 days post-fertilization (Dpf) for eye measurements, lens apoptosis detection, and cardiac looping assays, and juvenile fry were examined at 3 months postfertilization (Mpf) for effects on eye development. B. The effects of SAG on the expression of Shh pathway *gli1*, *ptch1*, and *nkx2.1* genes determined by qRT-PCR at the 13-15 somite stage. Asterisks indicate significance at p < .0001, p < .0001, and p = .001094 from left to right. Statistics by one-way ANOVA with Tukey HSP. C-I. Effects of SAG on eye development at 2.5 Dpf (C, D, G-I) and 3 Mpf (H, I). C-D. Lens apoptosis in SAG treated (D) but not control embryos (C). Arrows: lens. G-I. Reduction of the ventral optic cup in SAG treated (F) but not control (E) embryos. Eyes and lenses are outlined by white dashed lines. Scale bar in E is 100 µm; magnification is the same in E and F. G. Box and whisker plots showing eye size reduction by SAG. Horizontal lines: Medians. Vertical lines: whiskers representing the 25^th^ percentile. Asterisk indicates significance at p = .00052. N = 14. Statistics by one-way ANOVA with Tukey HSD. H, I. Eye degeneration in fry developed from SAG treated (F) but not control (E) embryos. J-L. Effects of SAG on KV size. KV size is increased in SAG treated (K) compared to control (J) embryos at the 13-15 somite stage. Arrows: KVs. Scale bar in J is 30 µm; magnification is the same in J and K. L. Box and whisker plots showing increased KV size in SAG compared to control embryos. Horizontal lines and vertical lines are as described above Asterisk; significance at p = .000031. N = 15. Statistics by one-way ANOVA with Tukey HSD. M-R. Effects of SAG on KV ciliation. KVs from control (M) and SAG (N) SF-D embryos stained with alpha-acetylated tubulin antibody at the 15 somite stage. Scale bar in M: 10 µm; magnification is the same in M and N. O-R. Box plots showing the quantification of KV cilia number (O), area (P), density (Q), and length (R) in 15 somite stage control and SAG treated embryos. N = 5 (control) and 7 (SAG). Asterisks: Significance in Q, p = .036914, and R, p =.001206. Statistics by one-way ANOVA with Tukey HSD. S. Effects of SAG on cardiac looping in SF-D embryos. N is shown on the right of each bar. Asterisk: significance at p = .000015. Statistics by Chi^2^ test (Chi^2^ statistic = 22.2448).

The SAG-treated SF-D embryos showed larger KVs compared to controls at the 13-15 somite stage (Fig. 5J-L), including a small subset with multiple KVs. Furthermore, SAG treatment showed KV ciliation similar to cavefish, including an increase in the length and decrease in the density of KV cilia compared to controls (Figs. 2; 5M-R). Lastly, increased levels of cardiac L-looping were seen in SAG-treated SF-D embryos compared to controls (Fig. 5S). The results suggest that Shh overexpression causes KV modifications and reversals in the normal pattern of L-R heart asymmetry in surface fish.

### Left-right asymmetry of *dand5* expression is modified in the cavefish KV

L-R determination involves asymmetric expression of the Nodal antagonist Dand5, which is caused by degradation of *dand5* mRNA and subsequent activation of *spaw* expression on the left side of the LRO (Marques et al., 2004; Montague et al., 2018; Minegishi et al, 2021). To investigate L-R Dand5 asymmetry in *Astyanax, dand5* expression was compared in 13-15 somite SF-D, CF-D, and CF-L/D embryos (Fig. 6). In situ hybridization showed that *dand5* was expressed exclusively in the KV region at this stage of development (Fig. 6A, D, G). In SF-D embryos, stronger *dand5* staining was detected on the right side than on the left side of the KV (Fig. 6B, C), indicative of normal L-R *dand5* asymmetry (Montague et al, 2018). Most CF-D and some CF-L/D embryos also exhibited preferential *dand5* staining on the right side of the KV (Fig. 6 E, F, H, I). In contrast, *dand5* was expressed abnormally (approximately symmetric or reversed L-R asymmetry) in the KVs of a significant number of CF-L/D embryos (Fig. 6H-I). Tailbud qRT-PCR indicated that the overall expression of *dand5* was not significantly different in 13-15 somite cavefish and surface fish embryos (Fig. 6J). The results show that normal L-R *dand5* expression is equalized or reversed in the KVs of a significant number of CF-L/D embryos.

**Figure 6.**
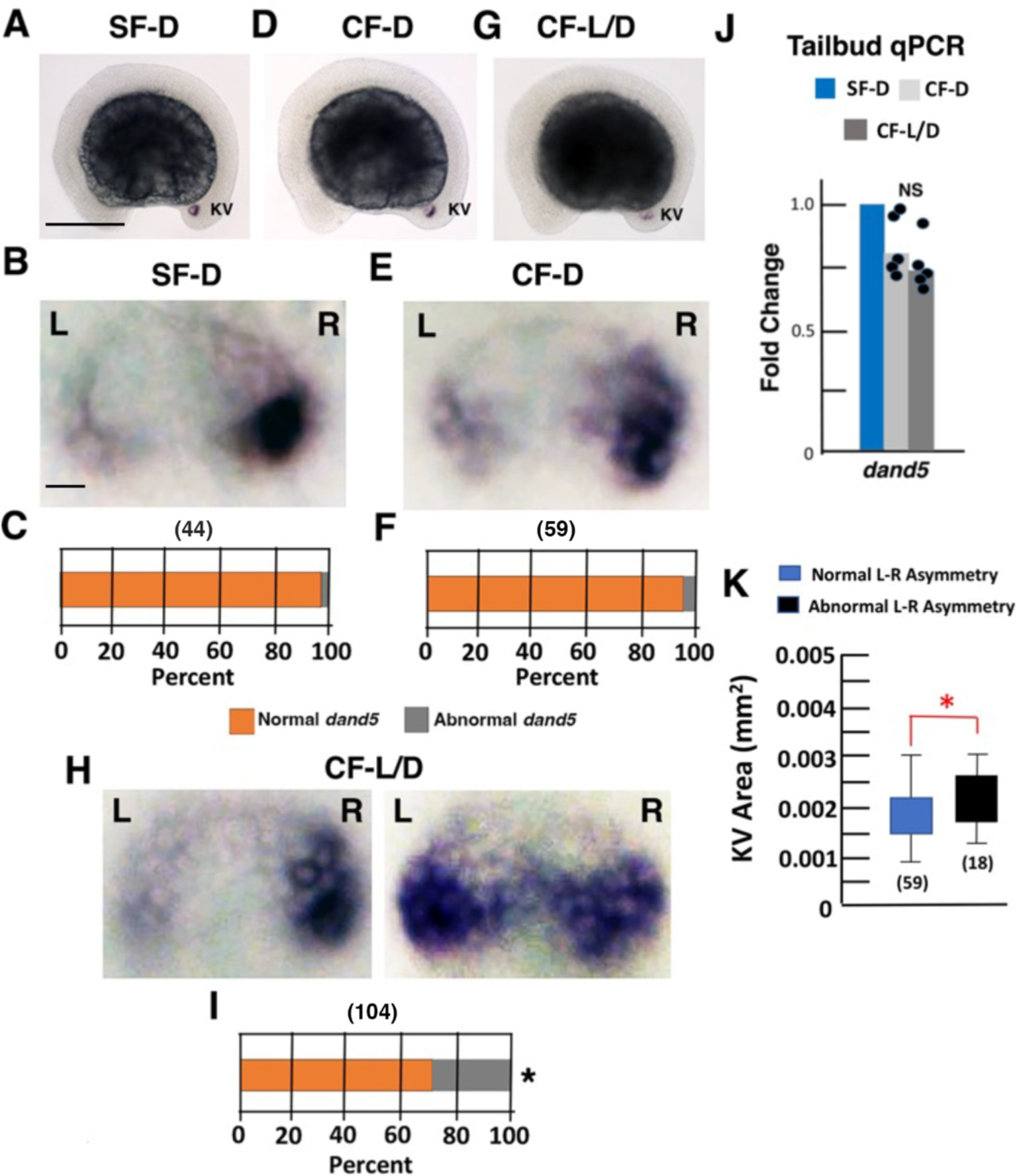
Left-right asymmetry of *dand5* expression is modified in the cavefish KV. A, B, D, E, G, H. In situ hybridization showing *dand5* expression in 13-15 somite SF-D (A, B), CF-D (C, D), and CF-L/D (G, H) embryos. Most SF-D (B) and CF-D (E) embryos and some CF-L/D (H, left) embryos show normal levels of *dand5* L-R asymmetry with stronger staining on the right compared to the left side of the KV, whereas other CF-L/D embryos (H, I) show abnormal *dand5* L-R asymmetry with similar levels staining on the right and left sides of the KV or more staining on the left than on the right side of the KV (not shown). L: Left. R: Right. Scale bar in A: 100 µM; same magnification in A, D, G. C, E, F. Scale bar in B: 10 µM: same magnification in B, E, H. Bar graphs showing the percentage of SF-D (C), CF-D (E), and CF-L/D (F) embryos with normal or abnormal *dand5* L-R asymmetry. The number of embryos analyzed is shown in parentheses on the top of each bar. Asterisk in G indicates significance at p < .00001. Statistics by Chi^2^ test comparing data in B, D, and G. Chi^2^ statistic = 25.9992. H. Tailbud qRT-PCR quantification of *dand5* mRNA levels in 13-15 somite SF-D, CF-D, and CF-L/D embryos. NS: no significance (p = .41066). Statistics by one-way ANOVA with Tukey HSD. K. Bar graphs showing the relationship between *dand5* asymmetry and KV size in 13-15 somite CF-L/D embryos. Asterisk: significance at p = .007395. Statistics by one-way ANOVA with Tukey HSD. Number of embryos analyzed is shown in parentheses at the base of each bar.

To determine if LRO size and changes in *dand5* asymmetry are related, we compared KV sizes in 13-15 somite CF-L/D embryos with normal and abnormal L-R *dand5* asymmetry. The results showed that embryos with larger KVs tended to have abnormal L-R *dand5* asymmetry, whereas embryos with smaller KVs showed normal L-R *dand5* asymmetry (Fig. 6J). These results suggest that development of a larger KV increases the likelihood of changing the normal pattern of L-R asymmetry in CF-L/D embryos.

### Shh overexpression affects left-right *dand5* asymmetry and Nodal laterality

To determine whether Shh signaling is involved in changing *dand5* L-R asymmetry, SF-D embryos were treated with SAG and the effects on asymmetric *dand5* expression in the KV were determined relative to controls (Fig. 7A-E). In situ hybridization showed that SAG-treated SF-D embryos developed significantly higher levels of abnormal *dand5* staining (approximately equal on each side of the KV or elevated on the left rather than the right side of the KV), which resembled modified *dand5* expression in CF-L/D embryos (Fig. 6H, I), compared to normal asymmetric staining in controls (Fig. 7A-D). Furthermore, tailbud qRT-PCR analysis indicated that SAG treatment failed to significantly change *dand5* expression levels in the KV region of SAG-treated SF-D embryos relative to controls (Fig. 7E, left), also similar to CF-L/D embryos (Fig. 6J). These results suggest that normal L-R *dand5* asymmetry is controlled by Shh signaling and that increases in the latter can affect *dand5* L-R asymmetry in the KV.

**Figure 7.**
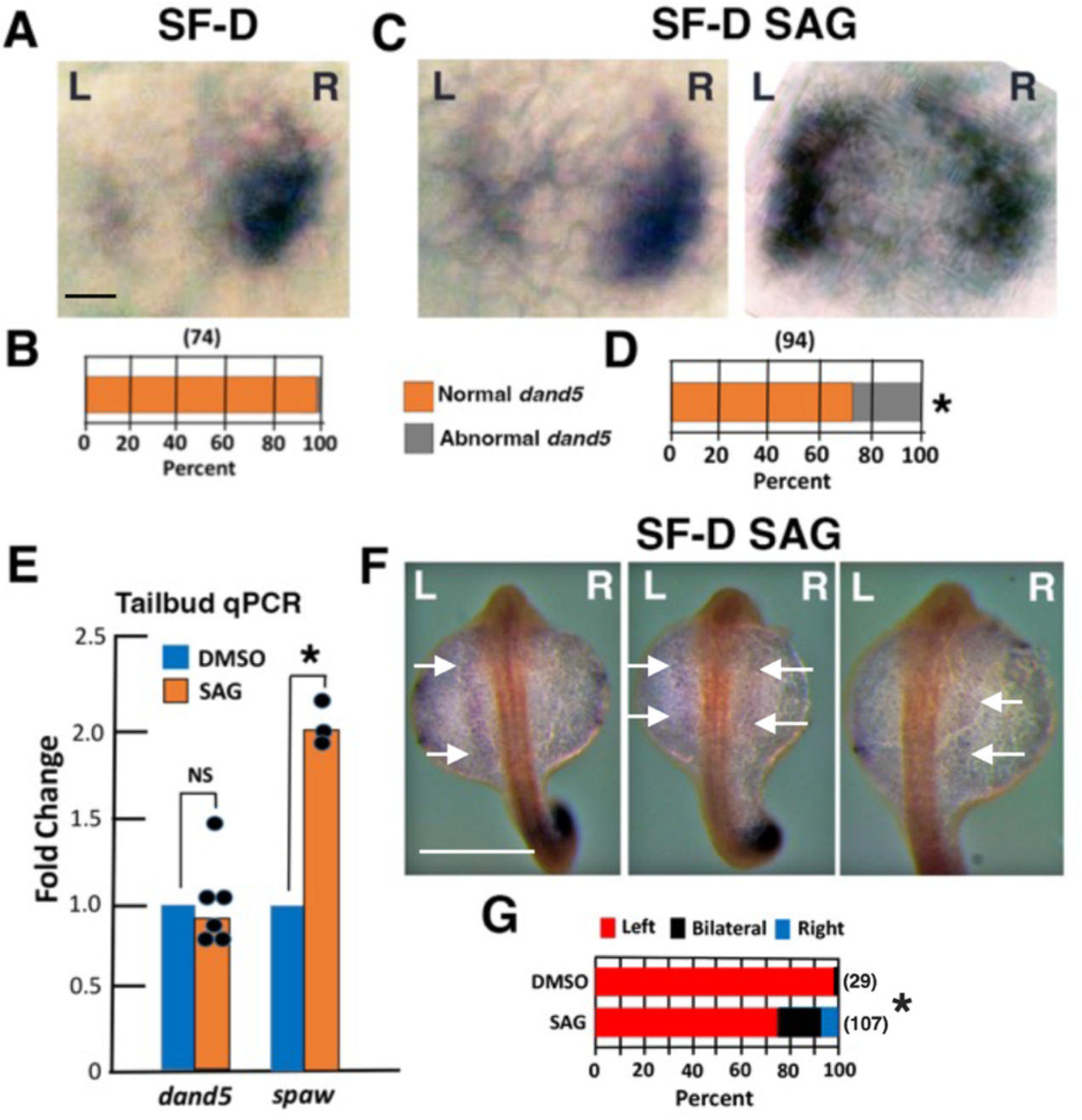
Sonic Hedgehog overexpression affects *dand5* Left-Right asymmetry and Nodal laterality in surface fish. A, C. In situ hybridization showing *dand5* expression in 13-15 somite control (A) and SAG-treated SF-D (C, D) embryos. Most control SF-D (A) and some SAG-treated SF-D (C, left) embryos show normal levels of *dand5* L-R asymmetry with stronger signal on the right compared to the left side of the KV, whereas other SAG-treated CF-L/D embryos (C, right) show abnormal *dand5* L-R asymmetry with similar staining on the right and left sides of the KV or more staining on the left than on the right side of the KV (not shown). L: Left. R: Right. Scale bar in A: 10 µM; same magnification in A, C, and D. B, D. Bar graphs showing the percentage of control (B) and SAG-treated (D) SF-D embryos with normal and abnormal *dand5* L-R asymmetry. The number of embryos analyzed is shown in parentheses on the top of each bar. Asterisk: significance at p =.057275. Statistics by a Chi^2^ test with data in B and D. F. Chi^2^ statistic = 3.61156. E. Tailbud qRT-PCR quantification of *dand5* (left) and *spaw* (left) mRNA levels in 13-15 somite control and SAG-treated SF-D embryos. NR: no significance. Asterisk: Significance at p <.00001 by one-way ANOVA with Tukey HSD. F. In situ hybridization showing *spaw* expression (arrows) the left lateral plate mesoderm (LPM) (left), both left and right LPM (middle), and right LPM (right) in 25-somite SAG-treated SF-D embryos. Scale bar in F (left) is 100 µm: magnification is the same in all frames. G. Bar graphs showing the percentage of 25 somite SF-D control (DMSO) and SAG-treated embryos with *spaw* expression in the left LPM, the both left and right LPM (bilateral, or the right LPM). Number of embryos analyzed is shown in parentheses on the right of each bar. Asterisk: significance at p = .035437. Statistics by Chi^2^ test. Chi^2^ statistic = 6.6243.

To address whether Shh overexpression affects Nodal signaling laterality in the LPM, *spaw* expression was examined in SAG-treated and control embryos (Fig. 7, E right-G). In situ hybridization showed that SF-D controls expressed *spaw* only in the left LPM, as described previously (Ma et al. 2021a), whereas *spaw* expression was detected in the left LPM, in the right LPM, or bilaterally in both the left and right LPMs, in SAG-treated 25-somite embryos (Fig. 7F, G). The number of SAG treated SF-D embryos with reversed or bilateral *spaw* expression was similar to that reported previously in CF-L/D embryos (Ma et al., 2021a). In addition, tailbud qPCR showed that *spaw* expression was significantly increased in SAG-treated compared to control SF-D embryos at the 13-15 somite stage, as would be expected from alteration of the normal pattern of *dand5* L-R asymmetry. We conclude that Shh overexpression affects the asymmetry of *dand5* expression in the KV, promoting changes in the laterality of Nodal signaling.

## Discussion

We have used the teleost *Astyanax mexicanus* to explore the mechanisms underlying natural changes in the highly-conserved pattern of L-R visceral organ asymmetry, which is difficult to study in other vertebrates. Features related to midline development and L-R asymmetry determination were compared in surface fish families exhibiting more than 95% normal left-oriented heart asymmetry (SF-D), cavefish families showing left-biased heart asymmetry resembling surface fish (CF-D), and cavefish families showing from 20-30% natural reversals of normal heart asymmetry (CF-L/D) (Ma et al., 2021a). The results revealed the effects of an enhanced Shh signaling center at the posterior end of the notochord on KV development and L-R asymmetry of *dand5* expression leading to the reversal of L-R heart laterality in cavefish embryos.

Previous studies have shown that enhanced Shh signaling in the pre-chordal plate at the anterior midline contributes to eye degeneration, increased olfactory and oral traits, and modified brain development in cavefish embryos (Yamamoto et al., 2004; 2009; Menuet et al., 2007; Pottin et al., 2011). Accordingly, upregulation of the Shh signaling system supports an adaptive explanation for the appearance of these regressive and constructive traits during cavefish evolution (Jeffery, 2005; Yamamoto et al., 2009), which is consistent with a selective sweep discovered in cavefish *shh* genes through genome sequencing (Moran et al., 2023). Here we extend this idea by showing that Shh signaling is also expanded at the posterior midline of cavefish embryos, and that posterior enhancement contributes to naturally-occurring changes in the pattern of cavefish heart asymmetry. Accordingly, we propose a revised model for midline effects in cavefish embryos compared to their surface fish counterparts that highlights enhanced Shh signaling at both the anterior and the posterior poles of the embryonic axis (Fig. 8). At the anterior midline, increased Shh signaling controls the development of optic primordia by modifying the expression of Pax6, Pax2, and Vax2 transcription factors (Yamamoto et al., 2004), whereas we show here that Shh upregulation at the posterior midline impacts the size and ciliation of the KV, L-R *dand5* asymmetry, downstream Nodal laterality, and the asymmetric pattern of heart development. In addition to Shh, the expression of other signaling systems, namely FGF, WNT, and BMP4, are also known to be modified in the anterior midline and contribute to evolutionary changes in cavefish embryos (Pottin et al., 2011; Hinaux et al. 2016; Ren et al., 2018). Likewise, we found that BMP4 expression also appears in the cavefish post-chordal plate, suggesting that multiple signaling pathways are active in a posterior center that is instrumental in mediating developmental changes between the two forms of *Astyanax*. The Shh-regulated *foxj1a* gene, which controls cilia development in the KV (Yu et al., 2008), is also overexpressed at the posterior midline of cavefish embryos where it is likely to be responsible for increases in the number of DFC, KV size, and the length of KV cilia. It will be interesting to determine whether variations in the strength of Shh signaling in the post-chordal region occur between different vertebrate species, and whether such variation could influence the evolution of traits that develop along the posterior midline, including those related to the regulation of L-R asymmetry.

**Figure 8.**
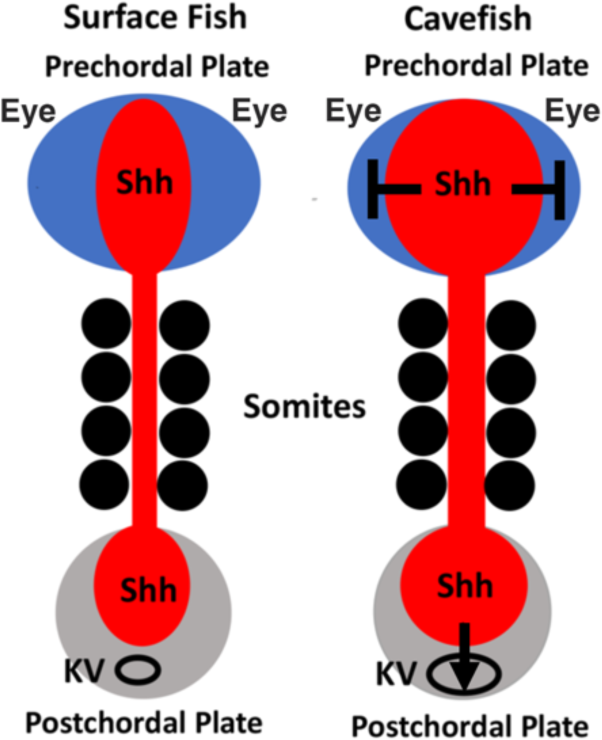
Diagram showing the model for Shh signaling in the surface fish and cavefish pre-chordal and post-chordal plates. Surface fish (left). Cavefish (right). Shh signaling is expanded in the cavefish pre-chordal plate near the regressing eyes and in the post-chordal plate near the enlarged KV. Shh expansion in the cavefish prechordal plate suppresses eye development and increases other cavefish traits, whereas Shh expansion in the post-chordal plate interferes with normal KV function in determining left-right organ asymmetry.

In this study, we discovered duplicated, or even triplicated, KVs, each aligned in parallel at the posterior end of the cavefish notochord. To our knowledge, multiple KVs or LROs have not been described in other teleost or vertebrate species. In *Astyanax*, multiple KVs were seen in about 10% of CF-L/D embryos, which show the highest levels of heart asymmetry reversal, and more rarely in CF-D embryos, but not in SF-D embryos. It is also notable that each of the multiple cavefish KVs was preceded by a posterior spur of the *shh*-expressing notochord, supporting a notochord contribution to DFC/KV induction, which has been proposed from other types of evidence in zebrafish embryos (Compagnon et al., 2014). The multiple KVs are probably a consequence of increases in the number and condensation foci of DFC, the KV precursors. The formation of double KVs in SAG-treated SF-D embryos, but not in controls, suggests that Shh overexpression is responsible for multiple KV development. However, the relatively low frequency of multiple KVs in CD-L/D embryos precluded an in depth investigation of their possible relationship to changes in L-R organ asymmetry. Future development of methods to increase the number of embryos with multiple KVs should allow their influence on L-R determination and Nodal laterality to be elucidated.

This study has revealed several changes related to enhanced posterior midline development and L-R asymmetry in cavefish. First, we found that DFC, the KV precursors, are more abundant in CF-L/D than in CF-D or SF-D embryos, and in some CF-L/D embryos DFC tend to aggregate into multiple foci, instead of a single focal point at the end of gastrulation, as in most CF-D and all SF-D embryos. DFC development is controlled by maternal factors in zebrafish (Moreno-Ayala et al., 2021), and if this is also the case in *Astyanax*, DFC (and KV) development could be targets of maternal effects responsible for the reversal of cavefish heart asymmetry (Ma et al., 2021a). Second, we found that CF-L/D KVs inflate to about twice the size of their CF-D or SF-D counterparts. There is a minimal size threshold for normal KV function in zebrafish embryos, and relatively modest increases in KV size do not change L-R laterality (Gokey et al., 2016). In contrast, our studies reveal that CF-L/D embryos with large KVs exhibit abnormal L-R *dand5* asymmetry (see below) and reversals in the pattern of heart asymmetry. Therefore, when KVs inflate to very large size, such as in CF-L/D embryos, there may indeed be downstream effects on visceral organ asymmetry. Third, we discovered two ciliary modifications in cavefish (both CF-L/D and CF-D) KVs compared to the surface fish KV: KV cilia are lengthened, which may be an effect of elevated expression of *foxj1a*, the master regulator of the ciliary developmental program (Yu et al., 2008; Stubbs et al., 2008), and cilia density is decreased in cavefish embryos. Thus, we propose that these changes could interfere with the counterclockwise ciliary beat responsible for generating leftward fluid flow in the KV and thus be responsible for changes in L-R asymmetry. Direct studies on ciliary movements will be needed to shed more light on the role of ciliary modifications in cavefish KV function.

Our results suggest that heart asymmetry reversal in cavefish is controlled at least in part by the enhancement of a Shh signaling center in the post-chordal plate, which influences the breaking of bilateral symmetry by the KV. The symmetry breaking process in teleosts and other vertebrates with ciliated LROs involves leftward ciliary beat and fluid flow in the KV, which leads to downregulation of the Nodal antagonist *dand5* on the left side of the KV and subsequent activation of *spaw* expression in the left LPM (Long et al., 2003; Hashimoto et al., 2004; Smith et al., 2021). We have shown here that *dand5* asymmetry is abnormal in some CF-L/D embryos, which show either approximately equal levels of *dand5* expression on both the left and right sides of the KV or more expression on the left than the right side of the KV, a pattern that was mimicked by Shh overexpression in SF-D embryos. These effects of Shh signaling on Dand5 asymmetry have not been described in any other vertebrate species.

However, it has been reported that Shh signaling mediates changes in L-R asymmetry by downregulating Dand5 in the cephalochordate amphioxus (Hu et al., 2017; Zhu et al., 2020). Together, our results and the amphioxus studies reveal a potentially important role for the ciliary-driven Shh-Dand5 axis in L-R symmetry breaking in the chordate phylum. At present, the mechanism of Shh suppression of *dand5* in cavefish is unknown. However, an intriguing possibility is that increased Shh signaling may affect the Bicc1-Ccr4 RNA complex that normally binds to and degrades *dand5* mRNA on the left side of the KV (Minegishi et al., 2021).

Our results also provide insight into how modifications in the conserved pattern of L-R organ asymmetry may have evolved in cavefish. Enhancement of Shh signaling increases the development of olfactory organs and oral structures involved in cavefish feeding (Menuet et al, 2007; Yamamoto et al., 2009; Hinaux et al., 2016). Moreover, Shh overexpression also contributes to the loss of eyes (Yamamoto et al., 2004), which may be an important factor in energy conservation (Moran et al., 2015). These constructive and regressive cavefish traits may be crucial for survival in the dark cave environment, where special challenges exist regarding sensory skills and energy budgeting. Since cavefish with normal or reversed heart asymmetry show no differences in survivorship (Ma et al., 2021a), there seems to be no advantage or disadvantage to changing L-R visceral symmetry. Therefore, reversed visceral asymmetry may have evolved in cavefish as a by-product of adaptive evolution related to the beneficial effects of Shh overexpression on traits subject to natural selection.

## Materials and Methods

### Biological materials

*A*styanax *mexicanus* surface fish and Pachón cavefish were raised in the laboratory in a constant flow aquatic system. We studied surface fish families showing >95% D cardiac looping (SF-D), cavefish families with >90% D cardiac looping (CF-D), and cavefish families with 20-30% L-cardiac looping (CF-L/D) (Ma et al., 2021a). The fish were raised in the laboratory at 25°C on a 14-h light and 10-h dark photoperiod. Spawning was induced by an increase in water temperature (Ma et al., 2021b), and embryos and larvae were raised at 23-25 °C. Experimental protocols were conducted according guidelines of the University of Maryland, College Park (IACUC #R-NOV-18-59; Project 1241065-1), and this study was carried out in compliance with ARRIVE guidelines.

### Heart asymmetry analysis

Heart asymmetry was determined at 2.5-3 days post-fertilization (dpf) in living larvae anesthetized with 2 mg/ml MS222 (Tricaine; Western Chemical Inc, Ferndale, CA, USA) or larvae fixed overnight with 4% paraformaldehyde (PFA) for immunostaining with the myosin heavy chain MF-20 antibody (Bader et al., 1982; Developmental Studies Hybridoma Bank, University of Iowa, Iowa City, IA, USA) using procedures described by Ma et al. (2021a) or visual inspection of cardiac tube looping from the ventral side under a Zeiss Axioskop compound microscope. Embryos were classified as having right (D) looping, no (straight) looping, or left (L) looping cardiac tubes.

### Kupffer’s vesicle analysis

The KVs of SF-D, CF-D, and CF-L/D embryos were viewed laterally or dorsal-posteriorly by live imaging under a Zeiss Axioskop microscope at the 10-12, 13-15, 16-18, and 19-21 somite stages, and KV areas were measured using ImageJ software. To quantify KV cilia, 13-16 somite stage embryos were fixed overnight in 4% PFA at room temperature to preserve microtubules, then dehydrated through increasing concentrations of methanol to 100%, and stored at -20°C. For cilia immunostaining, the rehydrated specimens were washed three times in PBST, blocked with bovine serum albumin and normal goat serum in PBST for 2 hours, stained with anti-acetylated tubulin antibody (1:500, cat# T7451, Sigma-Aldrich, St. Louis, MO, USA) at 4°C overnight, washed three times in PBST, and stained with Alexa Fluor-complex secondary IgG (ThermoFisher, Waltham, MA, USA). Z stack images of the immunostained KVs were photographed from their dorsal-posterior poles under a Zeiss LSM 980 Airyscan 2 laser scanning confocal microscope (Zeiss, Oberkochen, Germany). KV cilia were counted manually and lengths were measured using ImageJ.

### Dorsal forerunner cell staining and quantification

DFC were stained with the fluorescent dye SYTO 11 (cat# S7573, Invitrogen, Carlsberg, CA, USA) as described by Cooper and D’Amico (1996). SF-D, CF-D, and CF-L/D embryos were raised to the 60% epiboly stage, dechorionated by treatment with 0.2 mg/ml protease type XIV (cat# P5147, Sigma-Aldrich) for 1 min at room temperature, and then incubated for 30 minutes with SYTO 11 diluted in DMSO to a final concentration of 15 µM. Control embryos were treated with 2% DMSO in system water. At the 70%-80% epiboly stage, embryos were imaged and fluorescent DFC were quantified manually from photographic images.

### In situ hybridization

SF-D, CF-D, and CF-L/D embryos were dechorionated by protease treatment (see above) or manual removal with forceps, then fixed in 4% PFA overnight, dehydrated in a series of increasing methanol concentrations to 100%, and stored at−20 °C. RNA probes were prepared using oligonucleotide primers (Table S1) designed using sequence information from the *A*. *mexicanus* surface fish genome assembly (Warren et al., 2021) in the NCBI repository. In situ hybridization was carried out as described by Ma et al. (2014). After the completion of hybridization, the embryos were washed with PBST and incubated in BM Purple AP Substrate (Roche, Basel, Switzerland) at room temperature in the dark. After the signal developed, the reaction was terminated by rinsing the embryos in PBS. The embryos were processed through an increasing glycerol series in PBS and photographed using a Zeiss Axioskop compound microscope.

### Tailbud isolation and RNA extraction

SF-D, CF-D, and CF-L/D embryos were raised to the 13-15 somite stage and tailbuds were removed manually using Precision Glide metal needles (0.45mm x 16mm, BD Biosciences, Franklin Lakes, NJ, USA) under a stereomicroscope. The isolated tailbuds were immediately immersed in TRI Reagent Solution (Life Technologies, Grand Island, NY, USA) and total RNA was extracted using the Direct-zol RNA Microprep kit (Zymo Research, Irvine, CA, USA).

### Quantitative real time RT-PCR

Total RNA from whole embryos or isolated tailbuds prepared as described above was treated with RNase-free DNase I (Zymo Research) to remove genomic DNA. cDNA was synthesized using the SuperScript IV VILO Master Mix and oligo (dT)_20_primers (Sigma-Aldrich, St. Louis, MO, USA). Quantitative reverse transcriptase PCR (qRT-PCR) was performed using the TB Green Premix Ex Taq II (Takara Bio, San Jose, CA, USA) under the following cycling conditions: initial denaturation for 30 seconds at 95°C, then 40 cycles for 5 seconds at 94°C and 30 seconds at 60°C, and finally 5 sec at 95°C, 1 min at 60°C, and 95°C at the increments of 0.11°C/sec. The oligonucleotide primers (Table S2) for qPCR were designed using sequence information from the *A. mexicanus* surface fish genome assembly (Warren et al., 2021). The *rpl11* gene was used as a reference gene. The qPCR reactions were performed using a LightCycler 480 (Roche, Indianapolis, IN, USA). The ΔCt for each gene was calculated by subtracting the average Ct value of the reference gene. To compare gene expression, ΔΔCt was calculated by subtracting the average ΔCt of the control from the gene of interest. For graphical representation, the fold change was calculated as 2^−(ΔΔCt)^. The ΔΔCt values were used for statistical analyses.

### Shh overexpression in surface fish

Overexpression of the Shh pathway was carried out by treating surface fish embryos with the cell permeable Smoothened agonist SAG (cat# 11914, Cayman Chemical, Ann Arbor, MI, USA) (Stanton and Peng, 2010). A 15 mM SAG stock solution was prepared in DMSO. Beginning at about 50% epiboly, embryos were dechorionated as described above and treated with 1 µM SAG or DMSO (control) for about 8 hours until the 13-15 somite stage, then some embryos were processed for KV analysis as described above, and other embryos were processed for tailbud isolation, RNA extraction, and qRT-PCR as described above, or washed into fresh system water and raised until later stages of development. At 2-2.5 dpf, SAG treated embryos and controls were processed for eye size measurements and detection of lens apoptosis, as described below, and the direction of cardiac looping, as described above. In addition, some SAG and control DMO-treated embryos were raised to about 3 months post-fertilization to determine long-term effects of Shh overexpression.

### Eye measurement and lens apoptosis detection

SAG and DMSO treated surface fish embryos were raised until 2-2.5 dpf. Eye areas were measured by live imaging under a Zeiss Axioskop compound microscope using ImageJ software. Lens apoptosis was determined in the same SAG and control DMSO embryos as heart eye measurements were made by staining with 5 μg/ml Lysotracker Red DND 99 (Invitrogen) for 30 min in the dark as described previously (Ma et al., 2018). Stained larvae were anesthetized with MS222 (see above) and mounted on glass slides for imaging.

## Acknowledgements

We thank Ruby Dessiatoun, Karina LaCroix, Lauren Cox, and Barend Keller for animal care, Sophia DeMaria for assistance with tailbud surgery and RNA isolation, and Dr. Aniket V. Gore for early recognition of differences in KV size between cavefish and surface fish.

## Data Availability

All relevant data can be found within the article and its supplementary information.

## Supplementary Tables

**Table S1.**
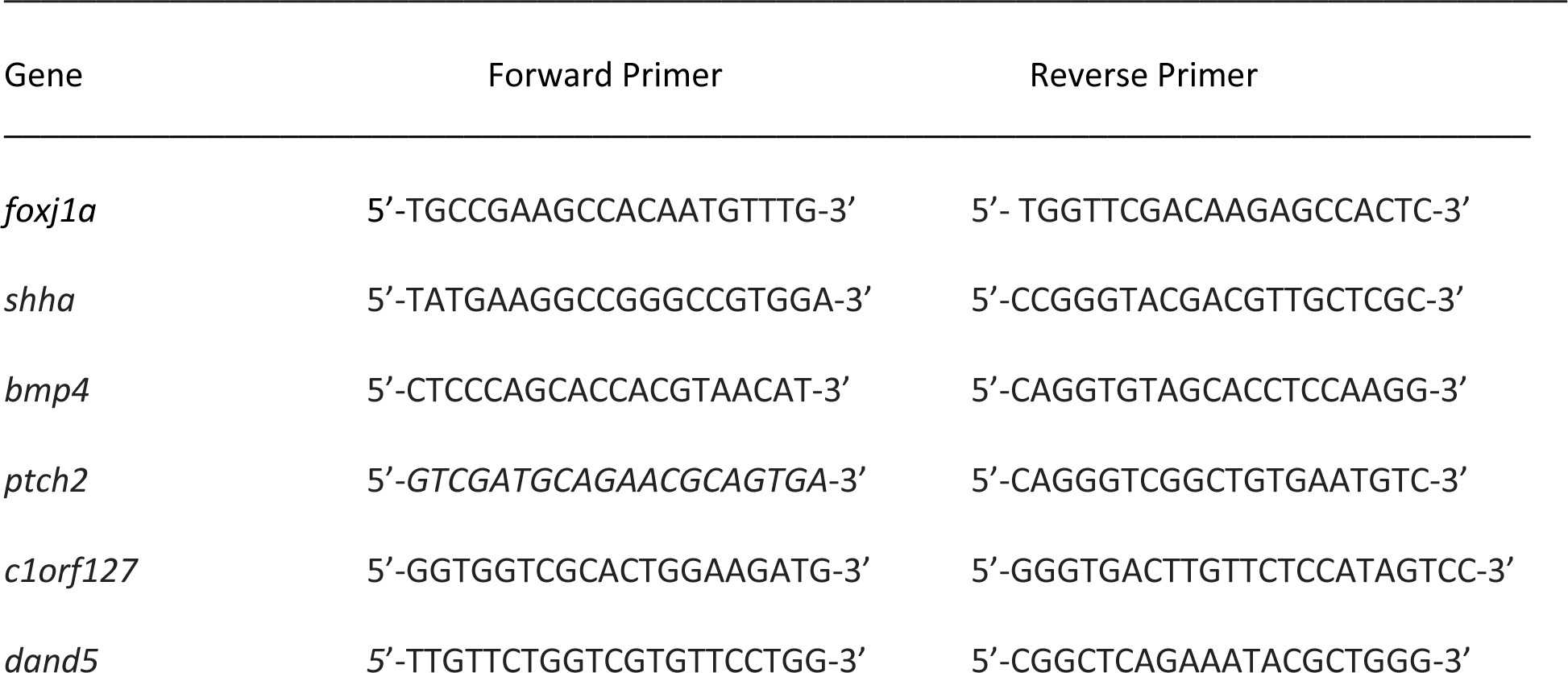
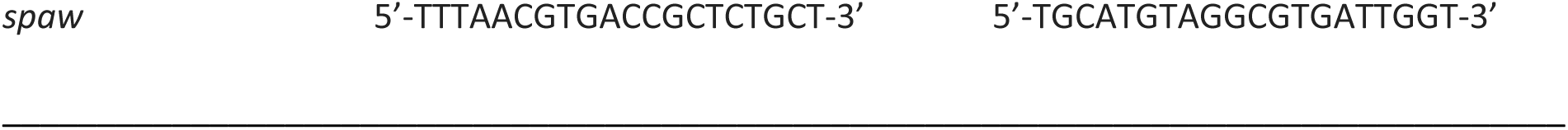
Primer sequences used for preparation of RNA probes used for in situ hybridization.

**Table S2.**
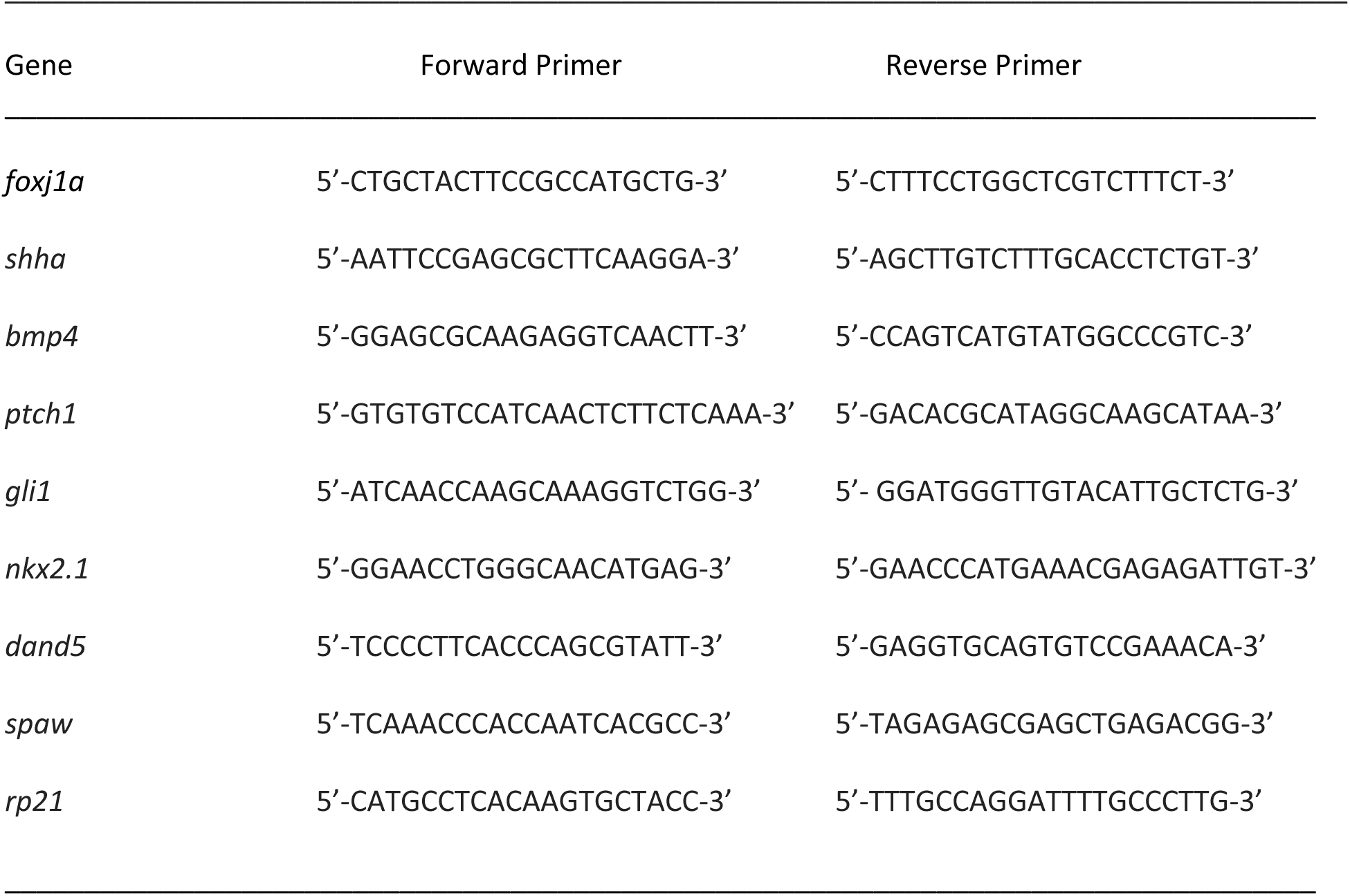
Primer sequences used for qPCR.

